# Positron Emission Tomography of CD47/SIRPα Axis and Image-Informed Therapeutic Design

**DOI:** 10.64898/2026.01.28.702416

**Authors:** Esther D. Need, Neetu Singh, Ayden Berndt, Alex Shelton, Samuel H. Cheshier, Shreya Goel, Sixiang Shi

## Abstract

CD47/SIRPα immune axis is of substantial clinical interest for innate cancer immunotherapy. Development on this axis has largely focused on monoclonal antibody agents and combination therapy strategies. Clinical use is challenging due to dose limiting side effects and severe anemia. Better understanding of the whole-body dynamics of CD47/SIRPα can be used to improve the developmental and therapeutic strategies targeting this axis. Herein, we developed anti-CD47 and anti-SIRPα radiotracers with good yields and stability. CD47/SIRPα biodistribution showed consistent whole-body results in healthy and colorectal cancer (CT26) allograft mice, demonstrating significant uptake in normal organs liver and spleen in addition to tumor accumulation of these agents. Enhancing immunogenicity via low-dose radiotherapy had no impact on over-all biodistribution but caused small, significant changes for anti-SIRPα tumor uptake. Antibody PEGylation of the anti-SIRPα tracer was further able to modify the whole-body distribution and reduce splenic uptake. These findings suggest that SIRPα targeted agents may benefit from co-therapies and drug delivery systems to optimize tumor uptake. Our work highlights the importance of in vivo molecular imaging in addition to in vitro and ex vivo assays when evaluating therapeutic designs.

## Introduction

Driven by exponential growth over the past decade, immunotherapy has evolved into the fourth pillar of cancer therapy. By blocking immune checkpoints that are overexpressed on cancer cells as a mechanism of immune escape, immune checkpoints inhibitors (ICIs) can effectively restore the body’s immune system to recognize and destroy cancer cells^1^. Great success has been achieved with ICIs for adaptive immune system; more than ten ICIs have been approved by the US Food and Drug Administration (FDA) targeting programmed cell death protein 1 (PD-1), programmed death-ligand 1 (PD-L1) and cytotoxic T-lymphocyte-associated protein 4 (CTLA-4), with more in clinical trials^2–4^. Discovered in the same era, the ICIs for innate immune system, although they are equally important and promising, have been developing at a much slower pace, leaving a significant gap in knowledge and underexplored niche of research and practices^5–7^. Besides the traditional measurement such as efficacy and side effects, we herein evaluate the ICIs for innate immune system via a unique perspective, the whole-body imaging with positron emission tomography (PET), which provides sensitive, quantitative, longitudinal and noninvasive delineation of their in vivo profiles.

CD47, famous as the “don’t eat me” signal, is one of the most important checkpoints for innate immune system. It is ubiquitously expressed on many tissues while highly expressed in many cancers as a mechanism of innate immune escape^8,9^. Its primary binding target, Signal Regulatory Protein alpha (SIRPα), is selectively expressed across myeloid cells, including macrophages and dendritic cells. When bound to CD47, SIRPα prevents critical cytoskeleton activation needed for phagocytosis, therefore playing an important role in immune tolerance and cancer immunotherapy^8,10^. The CD47/SIRPα axis also regulates tissue homeostasis such as self-tolerance^11^, pruning^12,13^, and red blood cell recycling^9,14^.

Cancers over express CD47 as a mechanism of innate immune escape^5,15–17^ and CD47 overexpression negatively correlates with patient outcomes^18^.Blocking CD47/SIRPα axis increases antibody-dependent cell-mediated cytotoxicity and phagocytosis, dendritic cell (DC) cross-priming, and natural killer (NK) cell cytotoxicity which can activate and enhance the adaptive immune response^19^. Therefore, targeting the CD47/SIRPα axis has emerged as a promising immunotherapy strategy, including monoclonal antibody ICIs, fusion proteins, bispecific antibodies, as well as small molecules, RNAis, and CAR-M therapies^20,21^. Among these therapeutic strategies, anti-CD47 monoclonal antibodies (mAbs) have dominated. Since the first clinical trial was halted due to a fatality^21,22^ anti-CD47 antibodies have greatly improved. Several clinical trials have demonstrated successful treatment with excellent efficacy and tolerable safety, such as Evorpacept (ALX148) in lymphoma and several types of solid cancers^23,24^. However, there are considerable side effects. The broad expression of CD47 can act as an antigenic “sink” potentially limiting therapeutic efficacy^25,26^. Specifically, erythrocyte and platelet expression of CD47 results in severe hemotoxic sideffects^19–21,27^. Interestingly, significantly fewer studies and clinical trials have been conducted on SIRPα-related therapy, the counterpart ligand of CD47 despite the rising argument that SIRPα-targeted immunotherapy can be safer than CD47-targeted immunotherapy. As such, the in vivo expression of SIRPα and biodistribution of SIRPα-targeted ICIs remain underexplored.

There has been continued interest in improving ICIs, including dosing strategies^19,28^, co-therapies^29–33^, developing tumor specific anti-CD47 antibodies^34^, and anti-SIRPα therapies^7,35^ in an attempt to mitigate these side effects. Recently, a few studies have investigated the dynamics of SIRPα expressing immune cells in vivo^36–38^. However, the effort to improve the whole-body physiological understanding of the CD47/SIRPα axis and how it may contribute to therapeutic challenges lags far behind the developmental work for therapies targeting this axis. Here, we analyzed the whole body biodistribution of the CD47/SIRPα axis by Zirconium-89 Positron Emission Tomography (PET) in healthy mice, a representative colorectal cancer allograft, a CT26 allograft post X-ray radiation therapy, and with a modified anti-SIRPα tracer. Our comprehensive descriptive biodistribution assessments can guide the development of CD47/SIRPα therapies to improve tumor targeting and response.

## Reagents and Methods

### Reagents

Cell culture and general supplies were purchased from Genesee. Chemicals were purchased from TCI chemicals (4’6-Diamidino-2-phenylindole dihydrochloride (DAPI), Ammonium Chloride NH^4^Cl), Sigma Aldrich (Sodium Carbonate (Na^2^CO^3^), potassium bicarbonate (KHCO^3^), Ethylenediaminetetraacetic acid disodium salt dihydrate (EDTA) 1M HEPES pH 7.0-7.6, Tween^®^ 20, and Chelex® 100 sodium form), and ThermoFisher (Paraformaldyde (PFA), 4% in PBS). Biologics were purchased from Sigma Aldrich (DNase I bovine pancreas and Bovine Serum Albumin (BSA), Worthington Biochem (Collagenase IV), and Jackson Immuno Research (Normal Mouse Serum). VECTASHIELD® Antifade Mounting Medium (H-1000-10) was purchased from Vector Laboratories and Instant Thin Layer Chromatography with silica gel (iTLC-SG) was purchased from Agilent.

Unmodified antibodies, including anti-mouse SIRPα (αSIRPα), anti-mouse CD47 (αCD47), and anti-trinitrophenol rat IgG2a isotype control (Isotype), were purchased from BioXcell. Conjugated flow cytometry antibodies were purchased from Biolegend except for Brillant Ultraviolet™ (BUV) 737 anti-MHC Class II purchased from Thermofisher. Amine-reactive reagents were purchased from Fisher Scientific (Alexa Fluor™ 488 Succinimidyl ester (AF488-NHS ester)), Lumiprobe (Cyanine 5 NHS ester), and Macrocyclics™ (p-SCN-Bn-Deferoxamine).

### Cell culture and in vitro assays

All cell lines were incubated at 37 °C with 5% CO^2^. CT26 Wild-type (WT), CT26 GFP-Luc, and 4T1 were culture in RPMI with 10% fetal bovine serum (FBS) and 1% penicillin-streptomycin (PS). MC38 and B16F10 were cultured in DMEM with 10% FBS and 1% PS. When cells reached 80% confluency, cells were sub-cultured by trypsinization, centrifugation, and re-seeding into TC treated flasks.

### Mice

Female 6-8 weeks old Balb/C (028BALB/C) and C57BL/6 (027C57BL/6) mice were purchased from Charles River. Tumor models were established by implanting 0.5 million cells (CT26 wild-type, CT26 GFP-Luc, MC38 or B16F10) or 0.2 million cells (4T1) under the skin of the right flank after shaving the area. Mice were monitored for body weight and tumor volume. When tumor volume reached 30 mm^2^, mice were used for flow cytometry, histology, or Positron Emission Tomography (PET) studies.

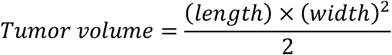

### Antibody radiotracer development

Antibodies (αSIRPα, αCD47, and IgG2a Isotype) were conjugated to p-SCN-Bn-deferoxamine (DFO) then labeling with Zirconium-89 (^89^Zr). All buffers used for radiotracer development were incubated with Chelex for at least 30 minutes prior to use.

For DFO conjugation, 0.6 mg of each antibody was diluted to 200 µL in PBS and DFO was added in a 10:1 ratio (40 nmol or 30 µg). The reaction was adjusted to pH 8.5-9.0 using 0.1 M Na^2^CO^3^, incubated for 2 hours at room temperature, then purified across a PD10 desalting column as before eluting with 0.5 M HEPES (pH 7). Products were confirmed by nanodrop UV-Vis spectroscopy.

400 µL of each DFO-antibody conjugate was radiolabeled with 37 MBq (1 mCi) ^89^Zr diluted into 400 µL of 0.5 M HEPES (pH 7). The reaction was confirmed to be pH 7 then left shaking for 1 hour at 37^°^C. ^89^Zr-DFO-antibody conjugates were purified by PD10 desalting column as before eluting with PBS (pH 7.4). Products were confirmed by activity yields.

### Radiotracer stability

Radiotracer stability was analyzed combining 50 µL of the ^89^Zr-DFO-antibody conjugates with 50 µL of either PBS or Mouse Serum. Assays were shaken at room temperature and sampled in triplicate by spotting 3 µL onto a radio iTLC plate after 1, 6, 24, 48, and 72 hours. iTLC plates were eluted in PBS, cut in half, and analyzed by gamma counting (Perkin Elmer 2270). The percent stability was calculated as below:

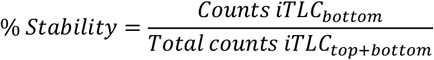

### Position emission tomography and ex-vivo biodistribution

Mice received 3.7 MBq (100 µCi) of ^89^Zr-DFO-αSIRPα, ^89^Zr-DFO-αCD47, or ^89^Zr-DFO-Isotype by tail vein injection. PET (Siemens Inveon MicroPET/CT) was performed at 6-, 24-, 48-, and 72-hours post injection. After the 72-hour PET scan, mice were scarified and tissues collected and weighed. Tissue radioactivity was recorded ex vivo by gamma counting (PerkinElmer).

Images were analyzed on Inveon Viewer, identifying regions of interest (ROIs) manually. All analyses were corrected for decay time and individual activity dose, reporting as percent injected dose per gram (%ID/g).

### Flow cytometry

Two-color flow cytometry (AF488-αCD47 or AF488-αSIRPα with APC anti-mouse CD45) was performed on tumor isolates from CT26WT, 4T1, MC38, and B16F10 allografts. Multicolor flow for innate immune cells was performed on tumor, spleen, and liver isolates from a CT26WT allograft. For multicolor flow, APC anti-mouse CD45, PE anti-mouse/human CD11b, PE/Cyanine 7 anti-mouse CD11c, PE/Cyanine 5 anti-mouse F4/80 (Biolegend), and BUV) 737 anti-MHC Class II (ThermoFisher) were used to sort immune cells for pan myeloid cells, macrophages, dendritic, and tissue cells.

αCD47 or αSIRPα (0.1 mg) were conjugated in house to AF488-NHS ester at a 20:1 molar ratio using the standard amine-NHS chemistry. The pH was adjusted to 7.5-8.0 using 0.1M KCO^3^. The reaction was protected from light and incubated for 2 hours at room temperature. AF488 antibody conjugates were purified across a PD10 desalting column eluting with PBS (pH 7.4), collecting the eluent from 2.8 to 3.8 mL. Antibody-dye yields and conjugation ratios were confirmed by nanodrop UV-Vis spectroscopy.

Mice were sacrificed and tumors, liver, and spleen tissues removed. Tumor and liver tissues were manually diced, digested with DNase (1 mg/mL) and Collagenase I (5 mg/mL) for 45 minutes at 37^°^C, then washed through a 70 µm filter with RPMI media to obtain single cells. Cells were centrifuged (300xg, 4^°^C, for 10 minutes), resuspended in 2-3 mL of erythrocyte lysis buffer (155mM NH^4^Cl, 10mM KHCO^3^, 500mM EDTA), and incubated for 5 minutes at room temperature. Lysis buffer was removed by centrifugation (300xg, 4^°^C, for 10 minutes). Spleen isolates were prepared in a parallel manner, but without enzymatic digestion and using a 40 µm filter Cells were resuspended in FACS buffer (2% FBS in PBS pH 7.4) at 2 million tumor cells per 100 µL or 1 million spleen or liver cells per 100 µL. Antibody-dyes were added and incubated for 1 hour at 4^°^C in the dark. Stained samples were washed twice and resuspended with FACS buffer and stained with DAPI (final concentration 3 µM). Two-color flow cytometry was performed in the same manner on tumor isolates for each tumor model.

Stained samples were stored on ice and data acquired immediately on a Cytoflex LX. Results were analyzed using FlowJo version v10.10 and GraphPad Prism 10.

### Immunofluorescent histology

Fluorescent antibody conjugates were prepared as for flow cytometry, using Cy5-NHS ester with each αCD47 or αSIRPα. Tumor, spleen, liver, and muscle tissues were harvested, frozen in O.C.T, and stored at - 80^°^C. Tissue slices (mm) were prepared by ARUP Research Histology Core Laboratory.

Tissue slides were fixed by submerging in cold PFA (4% in PBS) for 5 minutes and washing twice in PBS. The tissue regions were outlined with a hydrophobic PAP pen. In a dark humid environment, regions were blocked with 150 µL of 5% BSA, 0.1% Tween^®^ 20 in PBS at room temperature for 1 hour. The humidified chamber was transferred to a 4^°^C cold room, the blocking buffer was drained, and 150 µL of either Cy5-αCD47 or Cy5-αSIRPα at a concentration of 5 ug/mL was added and left staining overnight (16-18 hours). Tissue sections were drained, washed in PBS three times, and stained with 150 µL of 300 nM DAPI for 10 minutes at room temperature in the dark. Finally, slides were washed three times in PBS and cover slips were mounted using a drop (8-10 µL) of VECTASHIELD® Antifade Mounting Medium. Stained slides were stored dark at 4^°^C for no more than 3 days and imaged using a Leica SP8 Confocal microscope. CD47 images were acquired with a line sequence protocol with 5% power and 75% on the 405 nm laser and with 15% power and 40% gain on the 633 nm laser. SIRPα images were acquired with a line sequence protocol, with 5% power and 75% gain on the 405 nm laser and with 15% power and 75% gain on the 633 nm laser.

### X-Ray irradiation

Two days prior to PET imaging, tumor bearing mice were given a 10 Gy single fraction using a small animal radiation research platform (SARP, Xstrahl Inc.)

### Statistics

Statistical analysis was performed in GraphPad Prism 10 using a Mann-Whitney U test and reported when p<0.5.

## Results

### CD47/SIRPα imaging in healthy mice

PET imaging was conducted in healthy mice to visualize the expression of CD47 and SIPRα and the biodistribution of their targeted antibodies without any pathological influence. PET tracers were developed for anti-SIRPα antibody (^89^Zr-DFO-αSIRPα), anti-CD47 antibody (^89^Zr-DFO-αCD47) and an isotype control antibody (^89^Zr-DFO-Isotype). Antibodies were conjugated to Deferoxamine (DFO) via NHS chemistry and radiolabeling to Zirconium-89 as shown in Figure 1A. Tracers were purified with over 90% radiolabeling yields at 94 ± 1%, 92 ± 3%, and 95 ± 1% for ^89^Zr-DFO-αSIRPα, ^89^Zr-DFO-αCD47, and ^89^Zr-DFO-Isotype respectively. The tracer labeling remained stable after 92 hours in PBS at 98 ± 0.5%, 96 ± 1%, and 96 ± 2% and in mouse serum at 99 ± 0.3%, 97 ± 0.6%, and 97 ± 0.5% for ^89^Zr-DFO-αSIRPα, ^89^Zr-DFO-αCD47, and ^89^Zr-DFO-Isotype respectively (Figure 1B, C).

**Figure 1.**
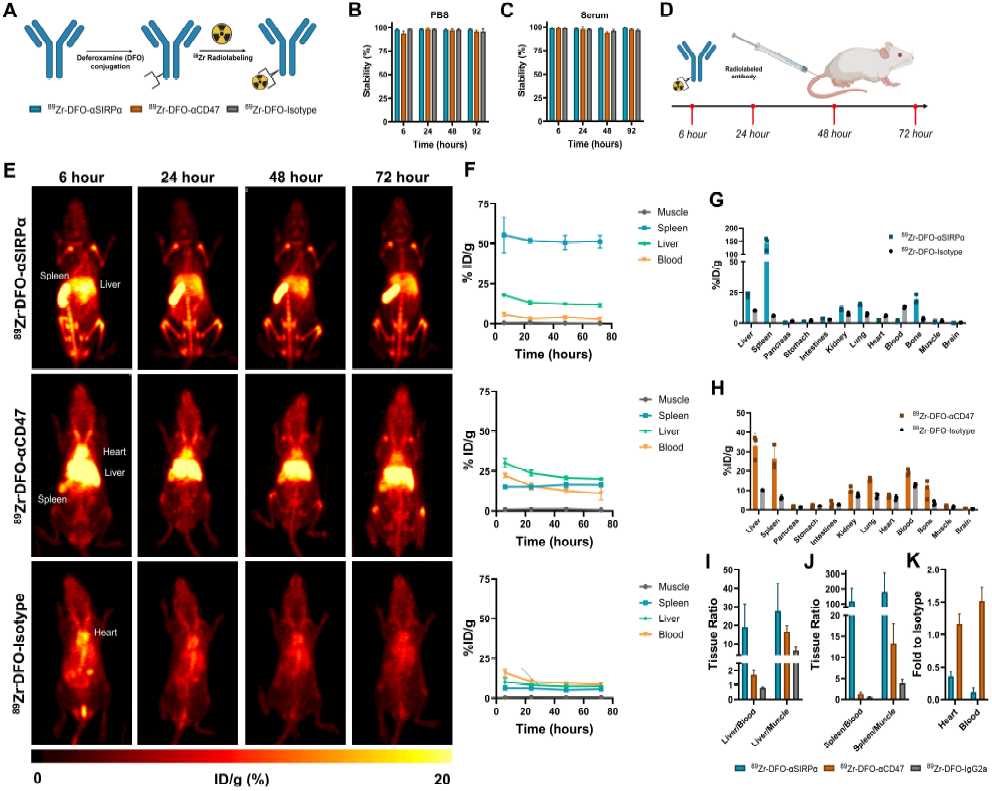
CD47/SIRPα biodistribution analysis in healthy mice. Conjugation and radiolabeling scheme for PET radiotracers (A) and their stability in PBS (B) and mouse serum (C) across the imaging time frame. PET imaging was performed at 6-24-48- and 72-post administration (D). Representative PET images (E) and the TACs (F) for the three radiotracers in healthy Balb/C mice. Gamma counting of post 72-hour terminal biodistribution of ^89^Zr-DFO-αSIRPα (G) and ^89^Zr-DFO-αCD47 (H) compared to the isotype tracer. Tissue ratios for liver (I) and spleen (J) and the fold accumulation compared to isotype (K) for each tracer. All data are presented as mean±s.d. * p<0.05.

The radiotracers (100 µCi) ^89^Zr-DFO-αSIRPα, ^89^Zr-DFO-αCD47, and ^89^Zr-DFO-Isotype were administered to healthy Balb/C mice by tail vein injection, PET was conducted across three days, followed by *ex vivo* biodistribution by gamma counting (Figure 1D). Compared to the isotype control (Figure 1E-H), ^89^Zr-DFO-αSIRPα showed extremely high accumulation in the spleen (142 ± 20% ID/g; Figure 1E, F) and liver (23 ± 2% ID/g; Figure 1F), as determined by gamma counting (Figure 1G). These spleen and liver results were supported by the tumor to blood and tumor to muscle signal ratios (Figure 1I, J). This finding suggests strong accumulation of SIRPα-positive immune cells in spleen and liver even at non-pathological conditions. Therefore, spleen and liver toxicity should be carefully considered in developing SIRPα-targeted ICIs. The clearance of ^89^Zr-DFO-αSIRPα was rapid, with the final blood signal ratio of 0.12-fold compared to isotype (Figure 1K). In contrast, ^89^Zr-DFO-αCD47 exhibited a more widely distributed pattern, where the spleen and liver retained 26 ± 5 and 32± 5% ID/g respectively (Figure 1H). Of note, there was elevated retention in the blood at 19 ± 1% ID/g (Figure1H), 1.5-fold that of the isotype control. This is consistent with the known CD47 expression on circulating red blood cells and immune cells under non-pathologic conditions.

### CD47/SIRPα imaging in CT26 allografts

To explore the behavior of the CD47/SIRPα axis in a cancer model and its role in therapy, four murine tumors were screened by flow cytometry for expression of CD47 on the tumor tissue cells (Figure 2A). Though all four mouse allografts showed slight enhancement in CD47 expression, CT26, a murine colorectal cancer, was selected for further work due to its 6.8-fold increase in CD47 expression compared to the isotype control (Figure 2B).

**Figure 2.**
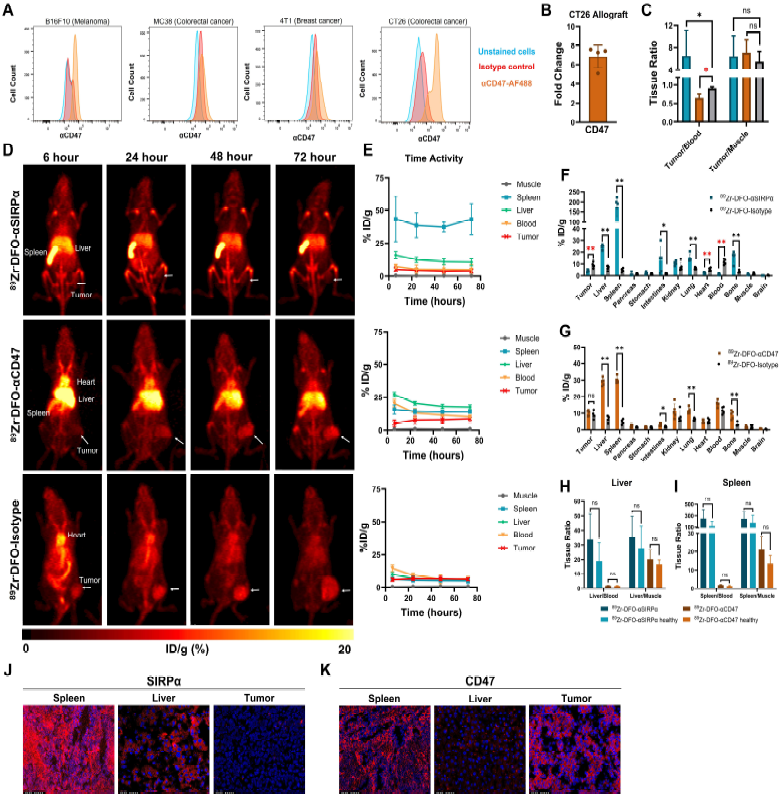
CD47/SIRPα biodistribution analysis in CT26 allografts. Representative histograms (A) from flow cytometry used to select a mouse allograft model for the highest CD47 expression (B) compared to isotype. Tumor to tissue ratios (C) from PET analysis with ^89^Zr-DFO-αSIRPα, ^89^Zr-DFO-αCD47, and ^89^Zr-DFO-Isotype. Representative PET images (D) and the time activity curves (TACs) (E). Gamma counting of terminal biodistribution for ^89^Zr-DFO-αSIRPα (F) and ^89^Zr-DFO-αCD47 (G) compared to the isotype tracer. Tissue ratios for liver (H) and spleen (I) for each tracer. Representative histology images for SIRPα (J) and CD47 (K) in spleen, liver, and tumor tissue. All data are presented as mean±s.d. * p<0.05.

PET revealed consistent whole-body distribution of the CD47/SIRPα axis between the CT26 allografts and healthy mice. ^89^Zr-DFO-αSIRPα showed high signal in the spleen (176 ± 42% ID/g) and liver (24.5 ± 1.8% ID/g) at 72 hours (Figure 2F), without any significant change when compared to the healthy mice (Figure 2H, I). Importantly, ^89^Zr-DFO-αSIRPα exhibited lower uptake in the tumor (4.2 ± 1.0% ID/g) compared to the isotype (9.0 ± 1.9% % ID/g) by ex vivo biodistribution (Figure 2D, E-F). Such low tumor uptake could have been caused by two possibilities: 1) CT26 tumors lacked sufficient infiltration of SIRPα-positive innate immune cells; 2) the extremely rapid and strong entrapment of αSIRPα in the spleen (176 ± 42% ID/g). The “sink-like” conditions created by the high-expression of SIRPα in the spleen deplete the circulating αSIRPα before it enters the tumor at substantial amounts. This phenomenon could explain why SIRPα-targeted ICI development has confronted major obstacles. Our findings through noninvasive PET imaging clearly highlight that low tumor uptake and strong splenic uptake are two key factors to be seriously considered in developing SIRPα-targeted ICIs. ^89^Zr-DFO-αCD47 accumulated at 29 ± 1% ID/g in the liver and 30 ± 2% ID/g in the spleen at 72 hours, much higher than the isotype control, indicating risk of liver and splenic toxicity for CD47-targeted ICIs. The tumor was visible with ^89^Zr-DFO-αCD47 reaching 10 ± 1% ID/g uptake at 72 hours. However, this was not statistically different from the isotype tumor accumulation as seen with ex vivo gamma counting (Figure 2D, E, G). The tumor-to-muscle ratio for each tracer ranged from 5.5 ± 1.5 to 7.1 ± 2 (Figure 2C). ^89^Zr-DFO-αCD47 showed a tumor-to-blood ratio of 0.65 ± 0.11 and ^89^Zr-DFO-αSIRPα had an elevated tumor-to-blood ratio of 6.5 ± 4.1, when compared to isotype (0.91 ± 0.03; p<0.05, Figure 2C). With promising tumor-to-blood accumulation, SIRPα was selected as the focus in our following studies, to explore if we can improve the *in vivo* profile of αSIRPα.

### Ex vivo validation of CD47/SIRPα expression

Immunofluorescence histology was performed on spleen, liver, and tumor tissue from CT26 allografts. SIRPα staining supported PET data with strong expression in the spleen, moderate expression in the liver, and minimal expression in the tumor tissue (Figure 2J). CD47 staining differed slightly from PET data with moderate expression on the spleen and tumor but minimal expression in the liver tissue (Figure 2K). This difference in the liver is likely due to the high blood retention of the αCD47 PET tracer and hepatic clearance observed in the in vivo study.

Multicolor flow cytometry was used to profile the cell types and expression of CD47/SIRPα in tumor, spleen and liver tissues and cells were gated according to supplemental figure 1. Tumor tissue had the highest expression of CD47, with a median fluorescent intensity (MFI) reaching 9.2 ± 1.5 × 10^5^ (Figure 3A, B). Additionally, nearly 100% of the tumor tissue cells showed over expression of CD47 while only 50-75% of the spleen and liver tissue cells highly expressed CD47 (Figure 3C). These results confirmed our original tumor profiling for CD47 expression. All immune cells showed elevated expression of CD47. Particularly, the tumor resident F4/80-myeloid cells and non-myeloid cells showed a statistically significant increase in CD47 expression compared to spleen and liver immune cells. One outlier was liver resident CD11c+ MHCII+ DCs, which had substantially higher expression of CD47 compared to the tumor and spleen resident DCs. The pattern of CD47 expression as determined by flow cytometry does exclude red blood cells which explains some of the variance from the PET data.

**Figure 3.**
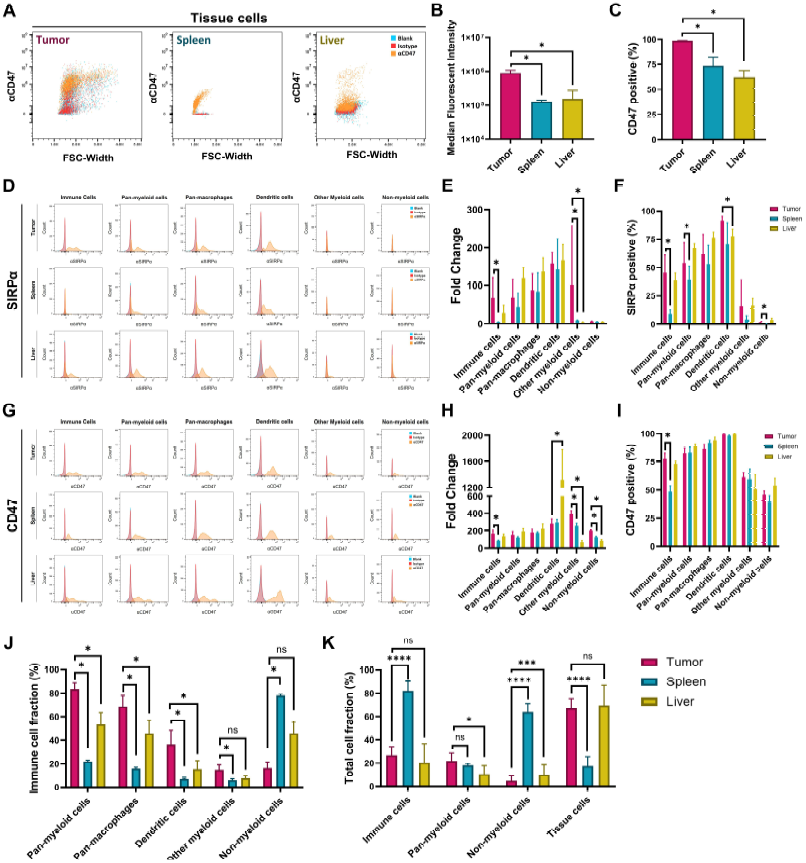
Multicolor flow analysis of SIRPα and CD47 expression from CT26 WT allografts. Representative dot plots for CD47 expression on CD45-tissue cells (A), their median fluorescent intensity (B), and percent of positive cells in the population (C). Representative histograms (D) for SIRPα expression on CD45+ immune cells and select subtypes with the fold change in SIRPα expression compared to isotype control (E) and the percent of positive cells in the population (F) from tumor, spleen and liver isolates. Representative histograms (G) for CD47 expression on CD45+ immune cells and select subtypes with the fold change in CD47 expression compared to isotype control (H) and the percent of positive cells in the population (I) from tumor, spleen and liver isolates. Tissue specific differences in the immune subtypes by percent (J) and the total immune infiltration (K). All data are presented as mean±s.d. * p<0.05.

All myeloid cells showed elevated expression of SIRPα, with CD11c+ MHCII+ DCs showing slightly higher fold expression than macrophages in their respective tissues. CD11b-non-myeloid cells showed minimal to no expression of SIRPα. There were minimal differences in the tissue-based expression of SIRPα, with other F4/80-myeloid cells being the only group with a significant difference in SIRPα expression. There was a higher fraction of high SIRPα expressing immune cells resident in the tumor compared to liver and spleen (Figure 3D, E and F) due to the abundance of lymphatic cells (non-myeloid cells) in liver and spleen which do not express SIRPα (Figure 3D).

Immune cell abundance and type varied dramatically from tissue to tissue. 84 ± 4% of the tumor resident immune cells were of the myeloid lineage, with macrophages composing 69 ± 8% and DCs 36 ± 10% of the immune cells. Non-myeloid lineage cells only composed 16 ± 4% of the tumor resident immune cells. Liver tissue had a close to a 50/50 immune lineage split between myeloid and non-myeloid, composing 46 ± 10% and 46 ± 8% respectively. The splenic immune population was primarily non-myeloid lineage, at 78 ± 1% (Figure 3J). Considering the total immune infiltration, 27 ± 7% of live cells in the tumor were of an immune lineage while 82± 8% of splenic cells were of an immune lineage. Myeloid lineage cells composed similar fractions of the total cell count in tumor, spleen, and liver tissues at 22 ± 7, 18 ± 2, and 11 ± 7% respectively (Figure 3K). The consistent expression of SIRPα on myeloid cells and the pattern of immune infiltration leads to the explanation of the observed PET data, with the immune cells simply not infiltrating the tumor at enough to cause tracer accumulation.

### Radiation therapy alters tumor accumulation of αSIRPα

Radiotherapy-induced immunogenicity has been thoroughly studied. It stimulates immune activation through various pathways and generation of proinflammatory cytokines, while also promoting the abscopal effect^30,39–41^. Additionally, several studies have indicated that radiotherapy alters CD47 expression on tumor cells suggesting it may be a critical combination method for SIRPα-targeted ICIs^19,40^. As low tumor uptake and high splenic uptake were major obstacles for SIRPα-targeted ICIs, as demonstrated by PET imaging, we employed low-dose X-ray irradiation to induce intratumoral immunogenicity and enhance infiltration of SIRPα-positive immune cells. CT26 allografts were treated with a single 10 Gy fraction two days prior to PET, examining the CD47/SIRPα axis (Figure 4A). The whole-body distribution of the CD47/SIRPα axis remained similar to untreated tumor-bearing mice and healthy mice, with strong spleen accumulation of ^89^Zr-DFO-αSIRPα and moderate liver and spleen accumulation of ^89^Zr-DFO-αCD47. No differences were seen in the tissue ratios for the spleen or liver (Figure 4I, J). Notably, there was a significant increase in total tumor accumulation of ^89^Zr-DFO-αSIRPα (8.2 ± 0.6% ID/g) when compared to untreated tumor bearing mice (4.2 ± 1.0% ID/g; Figure 4B). These ratios remained significant when comparing tumor/blood and tumor/muscle ratios (Figure 4C). The tumor signal remained low for ^89^Zr-DFO-αCD47 (12.8 ± 2.3% ID/g; Figure 4D, F, and H) and did not show a statistically significant change. Although the tumor uptake of ^89^Zr-DFO-αSIRPα was still much lower than spleen, the exciting increase of tumor uptake upon radiotherapy strongly suggests that additional intervention to stimulate immunogenicity (e.g. radiotherapy) is an effective method to arouse and synergize with SIRPα-targeted immunotherapy.

**Figure 4.**
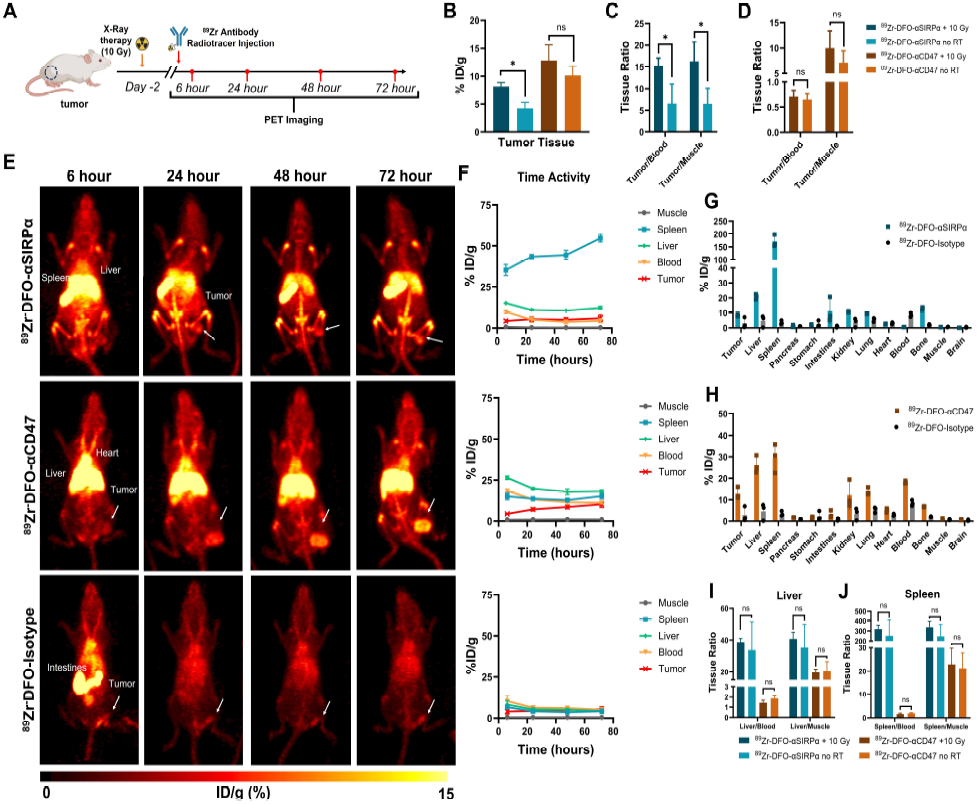
CD47/SIRPα biodistribution in CT26 allografts with single dose X-ray radiation therapy (A). Comparison of total tracer accumulation for tumor with and without radiation therapy (B). Tumor to tissue ratios for ^89^Zr-DFO-αSIRPα (C) and ^89^Zr-DFO-αCD47 (D). Representative PET images (E) and the time activity curves (TACs) (F). Gamma counting of terminal biodistribution for ^89^Zr-DFO-αSIRPα (G) and ^89^Zr-DFO-αCD47 (H) compared to the isotype tracer. Tissue ratios for liver (I) and spleen (J) comparing untreated and radiation treated mice.

### Antibody modification reduces splenic accumulation of αSIRPα

To explore the feasibility of further improving SIRPα targeting, we also tried to reduce its spleen uptake. αSIRPα was modified with a 5 kDa methyl-polyethylene glycol (mPEG) chain by NHS chemistry (Figure 5A), resulting in a PEGylated αSIRPα product (Figure 5B). αSIRPα-mPEG conjugations were performed in triplicate and consistently showed minimal reduction in SIRPα-mediated cell binding (Figure 5C). This modified αSIRPα-mPEG was further conjugated to DFO and labeled with ^89^Zr for PET. The ^89^Zr-DFO-αSIRPα-mPEG tracer showed over 95% stability in both PBS and mouse serum across 72 hours (Figure 5D, E). As before, 3.7 MBq (100 µCi) of the tracer was administered by tail vein injection to CT26 allografts which were then imaged across 72 hours (Figure 5F). There was a significant reduction in liver, spleen, lung, and bone joint accumulation of the tracer (Figure 5H). When normalizing to blood and muscle, the significant reduction in SIRPα accumulation remained, dropping from 251 ± 141 to 49 ± 18 (p<0.05) in spleen/blood ratio and from 246 ± 104 to 84 ± 18 (p<0.05) in spleen/muscle ratio (Figure 5J). There was no statistical difference in tumor accumulation at 72 hours (Figure 5H, K). However, there was a more rapid tumor accumulation of ^89^Zr-DFO-αSIRPα-mPEG showing a statistically higher tumor/blood ratio at 24 hours when the mPEG modified αSIRPα showed 2.7 ± 0.7 tumor/blood ratio compared to 1.1 ± 0.5 for the unmodified tracer (Figure 5L; p<0.5). This preliminary trial of antibody PEGylation is an excellent example of how antibody engineering can be used to reduce the splenic accumulation of αSIRPα and therefore enable functional SIRPα-targeted immunotherapy without severe splenic toxicity.

**Figure 5.**
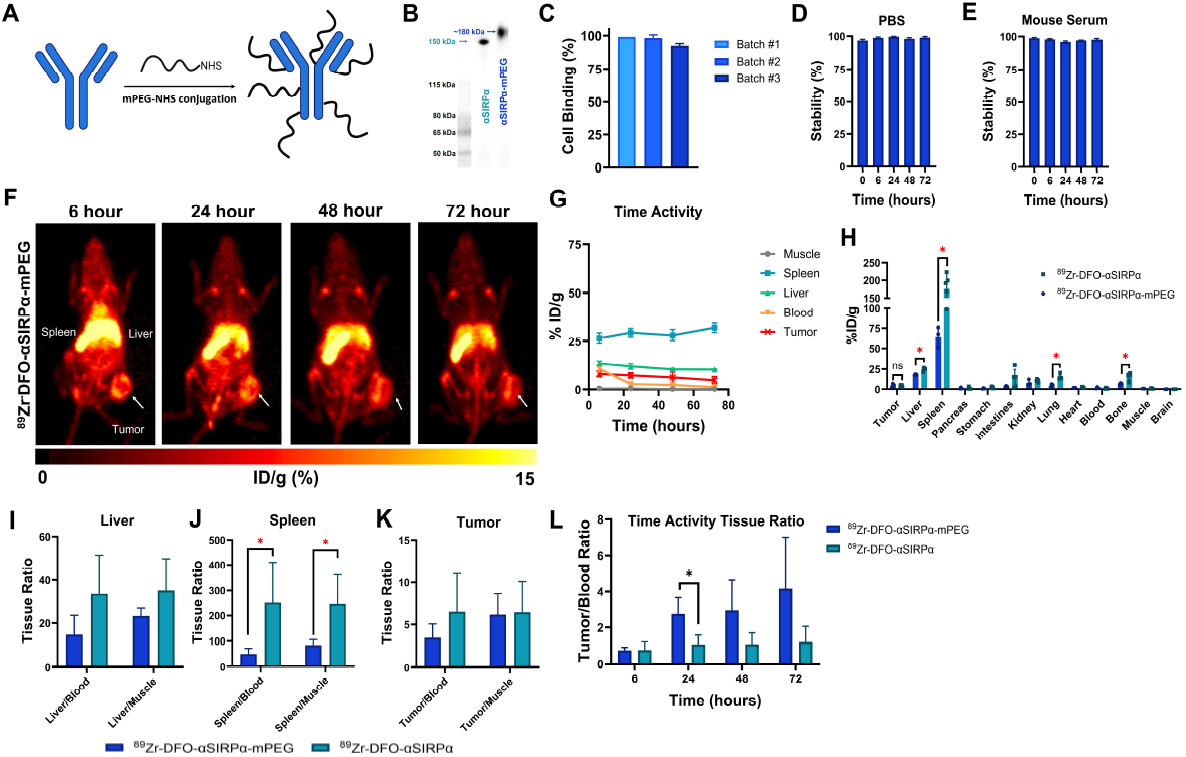
Altered SIRPα biodistribution with mPEG modification. mPEG modification of αSIRPα (A) and the size change shown by SAR-PAGE for 3 batches (B). Cell binding of αSIRPα-mPEG to SIRPα expressing RAW264.7 (C). Stability of ^89^Zr-DFO-αSIRPα-mPEG radiotracer in PBS (D) and mouse serum (E). Representative PET images (F) and the time activity curve (TAC) (G). Gamma counting of terminal biodistribution for ^89^Zr-DFO-αSIRPα-mPEG compared to previous unmodified ^89^Zr-DFO-αSIRPα (H). Liver (I), Spleen (J), and tumor (K) to tissue accumulation ratios comparing ^89^Zr-DFO-αSIRPα-mPEG and unmodified ^89^Zr-DFO-αSIRPα. Time analysis of tumor to blood ratio comparing ^89^Zr-DFO-αSIRPα-mPEG and unmodified ^89^Zr-DFO-αSIRPα (L). All data are presented as mean±s.d. * p<0.05.

## Discussion

Though there is high potential for innate immune modulation in cancer therapy by inhibiting the CD47/SIRPα axis, clinical application has been difficult with objective response rates (ORRs) under 5% for monotherapies in solid tumor pateints^19^. In response to the first drug trials being terminated due to hemolysis and anemia related toxicities^19,22^, most of the therapeutic development has focused on reducing toxicity with the development of RBC salvaging αCD47 antibodies and the weaker binding SIRPα-Fc fusion proteins. There have been 13 monoclonal αCD47 antibodies that have made it to clinical trials (Supplementary Table 1), 4 SIRPα-Fc fusions, and 1 small molecule.

Our results indicate that the sink hypothesis of CD47 is a legitimate challenge. In addition to peripheral blood, the liver and spleen retain 25-35% of the injected anti-CD47 tracer even in healthy mice. Combined with the near 20% retention in blood, these three compartments account for the majority of αCD47 binding at 72 hours (Figure 1H). Further, even in the CD47-expressing tumor model CT26, the liver and spleen retained similar amounts of the αCD47 tracer. The tumor accumulated 10 ± 1% ID/g, not significantly different from the tumor accumulation of the isotype control. The tumor/blood ratio of the αCD47 tracer was under 1 (0.65 ± 0.11). This opens the possibility that all tumor accumulation of αCD47 signal is the result of blood flow with no tumor specific accumulation. Histology showed higher tumor expression of CD47 than in the liver while PET showed significant liver signal, possibly attributed to the hepatic clearance of large therapeutics, including antibodies. The αCD47 agents now in clinical trials use RBC salvaging binding domains^21^, likely improving tumor-to-blood accumulation rates. Poor distribution can still impact therapeutic efficacy of CD47 blocking agents. A phase I study in cutaneous T-cell lymphomas saw 90% response with intralesional injections of TTI-621 a SIRPα-IgG1 Fc fusion protein (NCT02890368)^28^ much higher than the typical ORRs for hematological or solid cancers of 25.3% and 9.1% respectively^19^.

There have been very few attempts to therapeutically target SIRPα on the myeloid cells with only 2 SIRPα blocking agents in 3 clinical trials (Supplementary Table 2). However, there is some evidence that these agents are more effective in clinic^19^. Our study explained the challenges for SIRPα-targeted ICIs via a unique but important perspective, whole-body PET imaging. Our data indicates that the lack of benefit of SIRPα targeted agents is unlikely due to lower tissue expression or insufficient tumor infiltration of immune cells but related to the “sink” effect in the spleen. As SIRPα is primarily expressed on myeloid lineage cells (Figure 3) and the spleen retains 142±20% ID/g and 176±42% ID/g αSIRPα in healthy and tumor-bearing mice respectively (Figure 1G and Figure 2F), any gain from a lower “sink” in tissue or blood is lost in the dramatic uptake in immune organs.

There were several disagreements with PET biodistribution and the expression level of CD47 or SIRPα. For example, the flow cytometry and histology data suggested positive expression of CD47 in CT26 tumor and immune cells, whereas no elevation of αCD47 accumulation in tumor was detected by PET. In another example, the flow cytometry data suggested strong expression of SIRPα in tumor-associated myeloid immune cells, whereas tumor signal from αSIRPα was minimally detectable in PET. There are several reasons of these disagreements: 1) The flow cytometry provides relative expression of a specific biomarker while PET provides quantitatively accurate uptakes without influence of control groups, fluorescence compensations, and brightness of dyes. 2) The flow cytometry measures each organ in isolation, while PET images the whole-body pharmacokinetics, taking account of inter-organ competition where one organ may trap the tracer (i.e. the “sink” effect). 3) Flow cytometry overemphasizes the signal ratio but overlooks the quantity. However, the signal ratio may not matter if the quantity is at an extreme, too small or too large. For instance, only a very small portion of the cells in the spleen are SIRPα-positive myeloid cells. However, the spleen still has the highest uptake, because it is the largest immune organ in the body. These disagreements highlight the importance of PET imaging in developing ICIs, by providing more accurate, quantitative and longitudinal information of systemic pharmacokinetics and receptor engagement. Our work demonstrates that therapeutic development can be seriously misled by pure reliance on in vitro or ex vivo studies without validation of in vivo molecular imaging.

Combination therapy has much better success for both SIRPα and CD47 inhibition. Traditional radiotherapy and chemotherapies induce immunogenic cell death (ICD). ICD generates damage-associated molecular patters (DAMPs) which act as “find me”/”eat me” signals to the local immune cells^40,42^. CD47/SIRPα blockades in combination with radiation or chemotherapies are thought to reduce the “don’t eat me” signals while ICD generates an “eat me” signal, amplifying the phagocytic response synergistically^19,40^. Additionally, tumor associated CD47 expression has been shown to increase after both low-dose and ablative radiation therapy as a heightened manner of immune escape^29,39,43^. Despite the potential increase in tumor associated CD47, our results indicated that whole body sink challenges remain. Two days after a 10 Gy low dose radiation therapy, there was no change in anti-CD47 tracer accumulation in the tumor (Figure 4B, D). On the contrary, there was an increase in the tumor accumulation of the anti-SIRPα tracer (Figure 4B, C), aligning with the proinflammatory mechanism of radiotherapy^39^. These observations suggest that SIRPα may be a better target for combination therapies as has been seen in preclinical combination therapy studies^31^ and supported by clinical metanalysis indicating that anti-SIRPα combinations perform better (ORR of 28.3%) than anti-CD47 combinations (ORR of 3%)^19^. However, it is important to note that there was no significant change in the spleen or liver accumulation of either the anti-CD47 or anti-SIRPα tracers post radiation therapy (Figure 4I, J). “Sink” behavior could still present a challenge for dosing either ICI target.

Our PEGylated anti-SIRPα tracer had the same antibody binding affinity according to in vitro cell studies, while it showed a significant reduction of in vivo spleen “sink” accumulation (Figure 5). PEGylation showed no change to the toxicity of anti-SIRPα (Supplemental Figure 2). PEGylation is widely used to alter the biodistribution and pharmacokinetics of drug delivery systems. For proteins, PEGylation is primarily used to increase the serum half-life of small (less than 50 kDa) proteins^44^. PEGylation of nanoparticles and liposomes also increases the tumor accumulation of these drug formulations by the enhanced permeation and retention effect^45,46^. Neither mechanism can sufficiently explain the change in anti-SIRPα biodistribution due to the naturally long half-life of antibodies and the minor size change from 150 kDa to around 200 kDa compared to nanoparticles. One plausible explanation is that steric barrier created by PEG altered the binding kinetics of αSIRPα in the spleen, allowing for the antibody to circulate longer as evident by the higher blood uptake at early timepoints with 10.9±1.4%ID/g for the ^89^Zr-DFO-αSIRPα-mPEG compared to 7.0±1.4%ID/g for the ^89^Zr-DFO-αSIRPα tracer at 6 hours post injection (Figure 5G and Figure 3E). This would have contributed to the lower uptake in the spleen and higher uptake at early timepoints in the tumor for the ^89^Zr-DFO-αSIRPα-mPEG tracer (Figure 5H and 5L). However, the altered kinetics due to PEG also slightly reduce the binding kinetics of αSIRPα in tumor, which offset the increase of the antibody infiltration by 72 hours (Figure 5L). Further work is needed to understand the role of PEGylation on large molecular weight proteins such as antibodies. Regardless, this study demonstrates the potential of post-translational chemical modifications of antibodies to alter whole body biodistribution.

## Conclusion

Our work delineated the whole-body biodistribution profile of CD47/SIRPα axis via noninvasive PET imaging, demonstrating the importance of molecular imaging when developing therapeutics strategies targeting the CD47/SIRPα axis. Our data shows that the immune organs, especially the spleen and liver, strongly retain anti-CD47 and anti-SIRPα antibodies and act as a dose “sink” while minimal tumor uptake is acquired. To improve the tumor uptake and reduce the “sink” effect in the spleen, we conducted a series of experiments and demonstrated that radiotherapy-induced immune modulation can increase the tumor uptake of αSIRPα while PEGylation can significantly reduce its spleen uptake. This suggests radiation-based co-therapies and additional antibody modification can be beneficial for future ICIs targeting CD47/SIRPα axis.

## Acknowledgements

This work was supported, in part, by the University of Utah School of Medicine (S.S), University of Utah College of Pharmacy (S.G.), the University of Utah Immunology, Inflammation and Infectious Diseases (3i) Initiative (S.G. and S.S.), and the 5 For the Fight Fellowship (S.G). We acknowledge direct financial support from the Huntsman Cancer Center supported by the National Cancer Institute of the NIH under award number P30CA042014. The content is solely the responsibility of the authors and does not necessarily represent the official views of the NIH.

## References

1. Pardoll DM. The blockade of immune checkpoints in cancer immunotherapy. Nat Rev Cancer. 2012;12(4):252–264. doi:10.1038/nrc3239

2. Wang D, Bauersachs J, Berliner D. Immune Checkpoint Inhibitor Associated Myocarditis and Cardiomyopathy: A Translational Review. Biology. 2023;12(3). doi:10.3390/biology12030472

3. Cai JX, Wang SY, Hu H, et al. Disparities in the access to immune checkpoint inhibitors approved in the United States, the European Union and mainland China: a serial cross-sectional study. bmjph. 2025;3(1). doi:10.1136/bmjph-2024-001995

4. Twomey JD, Zhang B. Cancer Immunotherapy Update: FDA-Approved Checkpoint Inhibitors and Companion Diagnostics. AAPS J. 2021;23(2):39. doi:10.1208/s12248-021-00574-0

5. Kubli SP, Berger T, Araujo DV, Siu LL, Mak TW. Beyond immune checkpoint blockade: emerging immunological strategies. Nat Rev Drug Discov. 2021;20(12):899–919. doi:10.1038/s41573-021-00155-y

6. Wei G, Zhang H, Zhao H, et al. Emerging immune checkpoints in the tumor microenvironment: Implications for cancer immunotherapy. Cancer Letters. 2021;511:68–76. doi:10.1016/j.canlet.2021.04.021

7. Gauttier V, Pengam S, Durand J, et al. Abstract 1684: Selective SIRPa blockade potentiates dendritic cell antigen cross-presentation and triggers memory T-cell antitumor responses. Cancer Res. 2018;78(13_Supplement):1684. doi:10.1158/1538-7445.AM2018-1684

8. Morrissey MA, Kern N, Vale RD. CD47 Ligation Repositions the Inhibitory Receptor SIRPA to Suppress Integrin Activation and Phagocytosis. Immunity. 2020;53(2):290-302.e6. doi:10.1016/j.immuni.2020.07.008

9. Khandelwal S, Van Rooijen N, Saxena RK. Reduced expression of CD47 during murine red blood cell (RBC) senescence and its role in RBC clearance from the circulation. Transfusion. 2007;47(9):1725–1732. doi:10.1111/j.1537-2995.2007.01348.x

10. Khalaji A, Yancheshmeh FB, Farham F, et al. Don’t eat me/eat me signals as a novel strategy in cancer immunotherapy. Heliyon. 2023;9(10). doi:10.1016/j.heliyon.2023.e20507

11. Li D yang, Xie S li, Wang G yi, Dang X wei. CD47 blockade alleviates acute rejection of allogeneic mouse liver transplantation by reducing ischemia/reperfusion injury. Biomedicine & Pharmacotherapy. 2020;123:109793. doi:10.1016/j.biopha.2019.109793

12. Umemori H, Sanes JR. Signal Regulatory Proteins (SIRPS) Are Secreted Presynaptic Organizing Molecules *. Journal of Biological Chemistry. 2008;283(49):34053–34061. doi:10.1074/jbc.M805729200

13. Adams S, van der Laan LJW, Vernon-Wilson E, et al. Signal-Regulatory Protein Is Selectively Expressed by Myeloid and Neuronal Cells. The Journal of Immunology. 1998;161(4):1853–1859. doi:10.4049/jimmunol.161.4.1853

14. Burger P, Hilarius-Stokman P, de Korte D, van den Berg TK, van Bruggen R. CD47 functions as a molecular switch for erythrocyte phagocytosis. Blood. 2012;119(23):5512–5521. doi:10.1182/blood-2011-10-386805

15. Willingham SB, Volkmer JP, Gentles AJ, et al. The CD47-signal regulatory protein alpha (SIRPa) interaction is a therapeutic target for human solid tumors. Proceedings of the National Academy of Sciences. 2012;109(17):6662–6667. doi:10.1073/pnas.1121623109

16. Majeti R, Chao MP, Alizadeh AA, et al. CD47 Is an Adverse Prognostic Factor and Therapeutic Antibody Target on Human Acute Myeloid Leukemia Stem Cells. Cell. 2009;138(2):286–299. doi:10.1016/j.cell.2009.05.045

17. Sugimura-Nagata A, Koshino A, Inoue S, et al. Expression and Prognostic Significance of CD47–SIRPA Macrophage Checkpoint Molecules in Colorectal Cancer. International Journal of Molecular Sciences. 2021;22(5):2690. doi:10.3390/ijms22052690

18. The CD47-signal regulatory protein alpha (SIRPa) interaction is a therapeutic target for human solid tumors | PNAS. Accessed November 3, 2025. https://www.pnas.org/doi/abs/10.1073/pnas.1121623109

19. Son J, Hsieh RCE, Lin HY, et al. Inhibition of the CD47-SIRPα axis for cancer therapy: A systematic review and meta-analysis of emerging clinical data. Front Immunol. 2022;13. doi:10.3389/fimmu.2022.1027235

20. Qu T, Li B, Wang Y. Targeting CD47/SIRPα as a therapeutic strategy, where we are and where we are headed. Biomark Res. 2022;10(1):20. doi:10.1186/s40364-022-00373-5

21. Bouwstra R, van Meerten T, Bremer E. CD47-SIRPα blocking-based immunotherapy: Current and prospective therapeutic strategies. Clinical and Translational Medicine. 2022;12(8):e943. doi:10.1002/ctm2.943

22. EudraCT Number 2016-004372-22 - Clinical trial results - EU Clinical Trials Register. Accessed August 12, 2025. https://www.clinicaltrialsregister.eu/ctr-search/trial/2016-004372-22/results

23. Kim TM, Lakhani NJ, Soumerai J, et al. Evorpacept plus rituximab for the treatment of relapsed or refractory non-Hodgkin lymphoma: results from the phase I ASPEN-01 study. Haematologica. 2025;110(9):2102–2112. doi:10.3324/haematol.2024.286208

24. Lakhani NJ, Chow LQM, Gainor JF, et al. Evorpacept alone and in combination with pembrolizumab or trastuzumab in patients with advanced solid tumours (ASPEN-01): a first-in-human, open-label,multicentre, phase 1 dose-escalation and dose-expansion study. The Lancet Oncology. 2021;22(12):1740–1751. doi:10.1016/S1470-2045(21)00584-2

25. Lau APY, Khavkine Binstock SS, Thu KL. CD47: The Next Frontier in Immune Checkpoint Blockade for Non-Small Cell Lung Cancer. Cancers. 2023;15(21):5229. doi:10.3390/cancers15215229

26. Matlung HL, Szilagyi K, Barclay NA, van den Berg TK. The CD47-SIRPα signaling axis as an innate immune checkpoint in cancer. Immunological Reviews. 2017;276(1):145–164. doi:10.1111/imr.12527

27. Liu M, Liu L, Song Y, Li W, Xu L. Targeting macrophages: a novel treatment strategy in solid tumors. Journal of Translational Medicine. 2022;20(1):586. doi:10.1186/s12967-022-03813-w

28. Querfeld C, Thompson JA, Taylor MH, et al. Intralesional TTI-621, a novel biologic targeting the innate immune checkpoint CD47, in patients with relapsed or refractory mycosis fungoides or Sézary syndrome: a multicentre, phase 1 study. The Lancet Haematology. 2021;8(11):e808–e817. doi:10.1016/S2352-3026(21)00271-4

29. Rostami E, Bakhshandeh M, Ghaffari-Nazari H, et al. Combining ablative radiotherapy and anti CD47 monoclonal antibody improves infiltration of immune cells in tumor microenvironments. PLOS ONE. 2022;17(8):e0273547. doi:10.1371/journal.pone.0273547

30. Nishiga Y, Drainas AP, Baron M, et al. Radiotherapy in combination with CD47 blockade elicits a macrophage-mediated abscopal effect. Nat Cancer. 2022;3(11):1351–1366. doi:10.1038/s43018-022-00456-0

31. Ji K, Zhang Y, Jiang S, et al. SIRPα blockade improves the antitumor immunity of radiotherapy in colorectal cancer. Cell Death Discov. 2023;9(1):1–12. doi:10.1038/s41420-023-01472-4

32. Hsieh RCE, Krishnan S, Wu RC, et al. ATR-mediated CD47 and PD-L1 up-regulation restricts radiotherapy-induced immune priming and abscopal responses in colorectal cancer. Science Immunology. 2022;7(72):eabl9330. doi:10.1126/sciimmunol.abl9330

33. Lian S, Xie R, Ye Y, et al. Dual blockage of both PD-L1 and CD47 enhances immunotherapy against circulating tumor cells. Sci Rep. 2019;9(1):4532. doi:10.1038/s41598-019-40241-1

34. Li Y, Liu J, Chen W, et al. A pH-dependent anti-CD47 antibody that selectively targets solid tumors and improves therapeutic efficacy and safety. J Hematol Oncol. 2023;16(1):2. doi:10.1186/s13045-023-01399-4

35. Gauttier V, Pengam S, Durand J, et al. Selective SIRPα blockade reverses tumor T cell exclusion and overcomes cancer immunotherapy resistance. J Clin Invest. 2020;130(11):6109–6123. doi:10.1172/JCI135528

36. Nimmagadda S. SIRPα Immuno-PET Provides a Pan-Myeloid View of Immune Activity During COVID-19. J Nucl Med. Published online November 13, 2025:jnumed.125.270781. doi:10.2967/jnumed.125.270781

37. Wagner TR, Blaess S, Leske IB, et al. Two birds with one stone: human SIRPα nanobodies for functional modulation and in vivo imaging of myeloid cells. Front Immunol. 2023;14:1264179. doi:10.3389/fimmu.2023.1264179

38. Stammes MA, Koopman G, Wagner TR, et al. Noninvasive Monitoring of Inflammatory Processes by Myeloid Cell–Directed PET Tracers in an Experimental Severe Acute Respiratory Syndrome Coronavirus 2 Infection Model. J Nucl Med. 2026;67(1):145–151. doi:10.2967/jnumed.125.269721

39. Ozpiskin OM, Zhang L, Li JJ. Immune targets in the tumor microenvironment treated by radiotherapy. Theranostics. 2019;9(5):1215–1231. doi:10.7150/thno.32648

40. Xi Y, Chen L, Tang J, Yu B, Shen W, Niu X. Amplifying “eat me signal” by immunogenic cell death for potentiating cancer immunotherapy. Immunological Reviews. 2024;321(1):94–114. doi:10.1111/imr.13251

41. Rauf S, Smirnova A, Chang A, Liu Y, Jiang Y. Immunogenic Cell Death: the Key to Unlocking the Potential for Combined Radiation and Immunotherapy. bioRxiv. Preprint posted online February 17, 2025:2025.02.14.638342. doi:10.1101/2025.02.14.638342

42. Xiao L, Zhang L, Guo C, et al. “Find Me” and “Eat Me” signals: tools to drive phagocytic processes for modulating antitumor immunity. Cancer Communications. 2024;44(7):791–832. doi:10.1002/cac2.12579

43. Menaa C, Fan M, Lu HC, et al. Abstract 4963: The dynamic change of CD47 expression promotes tumor burden, metastases and resistance of breast cancer cells to radiotherapy. Cancer Research. 2013;73(8_Supplement):4963. doi:10.1158/1538-7445.AM2013-4963

44. Turecek PL, Bossard MJ, Schoetens F, Ivens IA. PEGylation of Biopharmaceuticals: A Review of Chemistry and Nonclinical Safety Information of Approved Drugs. Journal of Pharmaceutical Sciences. 2016;105(2):460–475. doi:10.1016/j.xphs.2015.11.015

45. Khan DR, Webb MN, Cadotte TH, Gavette MN. Use of Targeted Liposome-based Chemotherapeutics to Treat Breast Cancer. Breast Cancer(Auckl). 2015;9s2:BCBCR.S29421. doi:10.4137/BCBCR.S29421

46. Zalba S, ten Hagen TLM, Burgui C, Garrido MJ. Stealth nanoparticles in oncology: Facing the PEG dilemma. Journal of Controlled Release. 2022;351:22–36. doi:10.1016/j.jconrel.2022.09.002

