## Supplemental Figures and Tables for "Positron Emission Tomography of CD47/SIRPα Axis and Image-Informed Therapeutic Design"

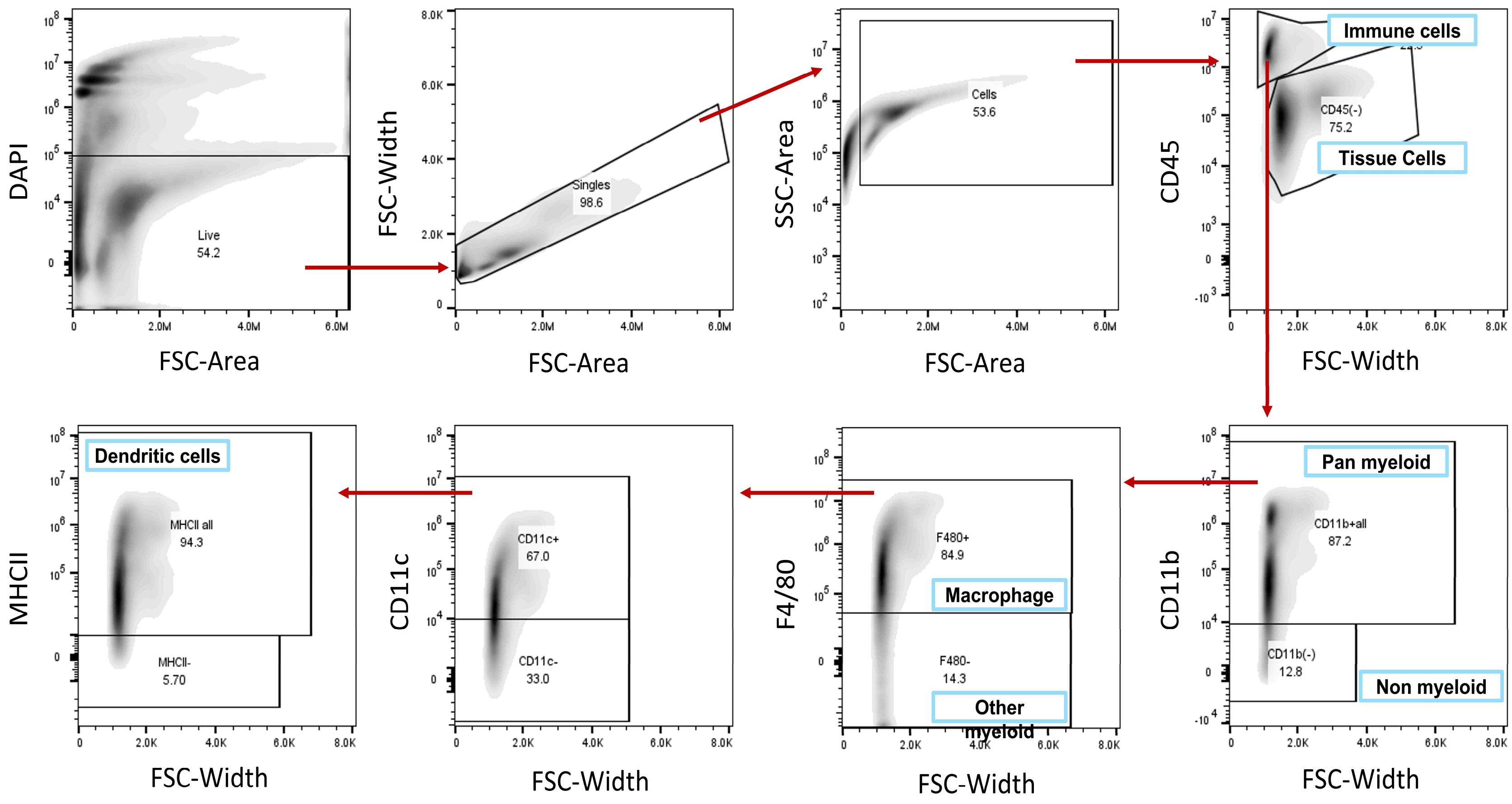


**Supplemental Figure 1** Multicolor flow gating strategy for tumor, liver, and spleen isolates separating tissue cells, immune cells, pan-myeloid, macrophage, dendritic cells, F4/80 negative myeloid cells, and CD11b- non-myeloid cells.

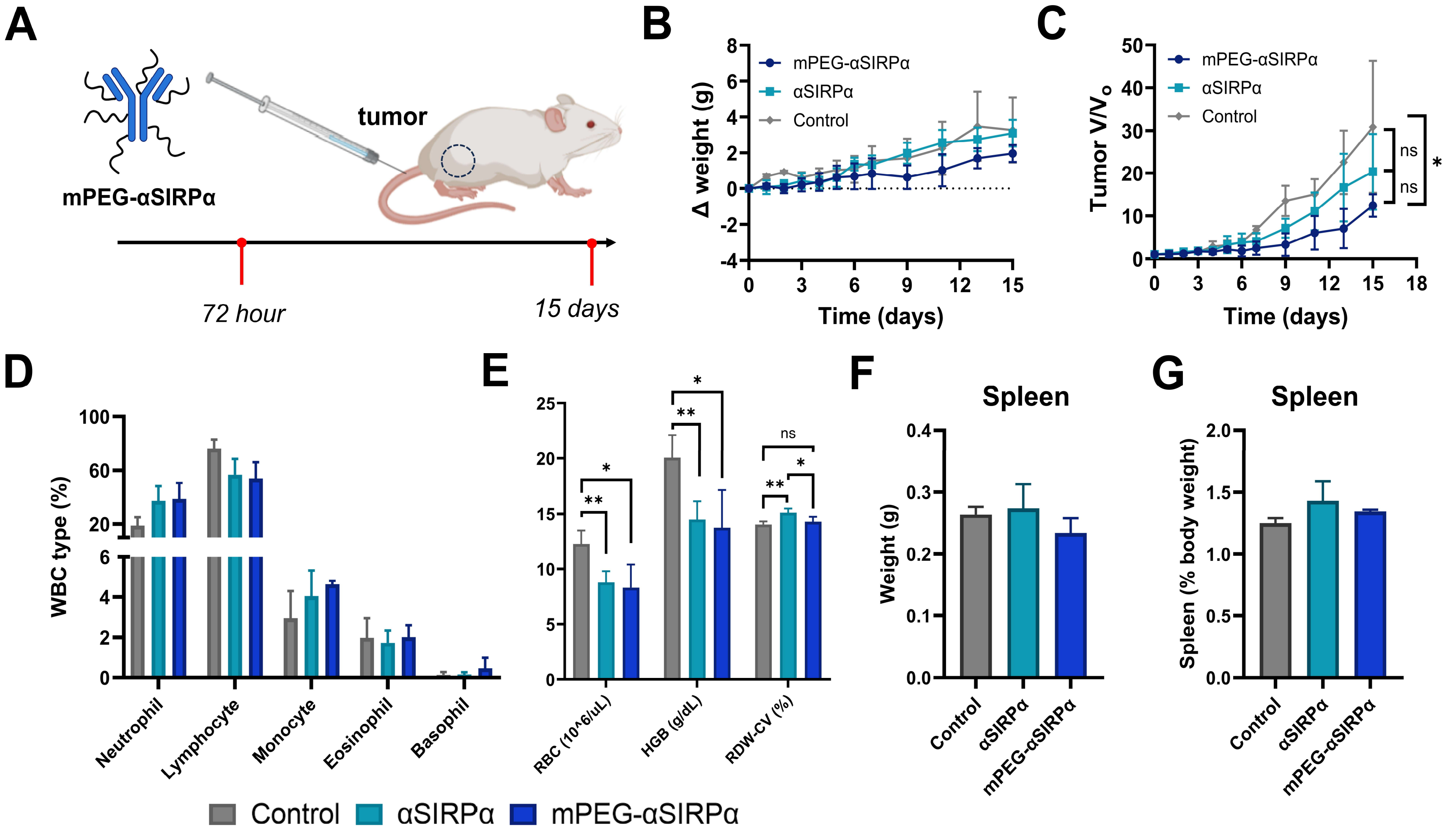


**Supplemental Figure 2** Toxicity profile of αSIRPα-mPEG at a single therapeutic 20mg/kg dose in CT26 allografts (A). Mouse monitoring for body weight (B) and change in tumor volume (C) across 15 days. Comparison of white blood cell sub types at 72 hours (D) and red blood cell parameters (E) at 72 hours. Change in total spleen weight (F) and as fraction of body weight (G) at 15 days post therapy. All data are presented as mean±s.d. * $p<0.05$.

**Supplementary Table 1** CD47 targeted agents in clinical trials for cancers

| Drug Format | Drug Name/Code | Clinical trial ID | Co-therapies | Cancer Types | Phase | N | Trial Status |
|---|---|---|---|---|---|---|---|
| Monoclonal anti-CD47 IgG2 | AO-176 | NCT03834948 | Paclitaxel, Pembrolizumab | Solid Tumor | I/II | 57 | Completed |
| | | NCT04445701 | Dexamethasone and Dexamethasone + Borezomib | Multiple Myeloma | I/II | 10 | Completed |
| Monoclonal anti-CD47 IgG4 | Letaplimab (IBI188) | NCT03763149 | NA | Advanced Malignancies | I | 20 | Completed |
| | | NCT04485052 | NA | Acute Myeloid Leukemia | I/II | 222 | Suspended |
| | | NCT04861948 | GM-CSF, Cisplatin/Carboplatin, Bevacizumab, Sintilimab, Pemetrexed | Solid Tumors, Lung Adenocarcinomas, and Osteosarcoma | I | 9 | Terminated |
| | | NCT04485065 | Azacitidine | Myelodysplastic Syndromes | I | 120 | Suspended |
| | | NCT04511975 | Azacitidine | Myelodysplastic Syndromes | I | 32 | Suspended |
| | | NCT03717103 | NA | Hematologic Neoplasms | I | 49 | Completed |
| | Ligufalimab (AK117) | NCT05229497 | Ivonescimab, Carboplatin, Cisplatin, and 5-Flourouracil | Solid tumor | Ib/II | 114 | Recruiting |
| | | NCT05214482 | AK112 and chemotherapy | Malignant Tumors | Ib/II | 250 | Active |
| | | NCT05235542 | AK104, Capecitabine, Oxaliplatin, Cisplatin, Paclitaxel, Ironetcan, Docetaxel, and 5-Fluruoracil | Advanced Malignant Tumors | Ib/II | 130 | Completed |
| | | NCT04900350 | Azacitidine | Myelodysplastic Syndrome | I/II | 136 | Active |
| | | NCT04980885 | Azacitidine | Acute Myeloid Leukemia | Ib/II | 68 | Active |
| | | NCT06508606 | Anti-EGFR | Head and Neck Squamous Cell Carcinoma | II | 10 | Not Yet Recruiting |
| | | NCT06387420 | Azactidine + Venetoclax | Acute Myeloid Leukemia | Ib/II | 180 | Recruiting |
| | | NCT05960955 | Cadonilimab, Oxaliplatin, Tegafur-gimeracil-oteracil potassium, Docetaxel, 5-Fluorouracil | Gastric and Gastroesophageal Junction Adenocarcinoma | II | 90 | Recruiting |
| | | NCT06789848 | Cadonilimab | Hepatocellular Carcinoma and Biliary Tract Cancer | II | 64 | Recruiting |
| | | NCT06601335 | AK112 and Pembrolizumab | Head and Neck Squamous Cell Carcinoma | III | 510 | Recruiting |
| | | NCT04349969 | AK104 | Malignant Neoplasms | I | 38 | Completed |
| | | NCT06953999 | Ivonescimab, Albumin -bound Paclitaxel, and Gemcitabine | Pancreatic Cancer | III | 999 | Not Yet Recruiting |

| | | | | | | |
|---|---|---|---|---|---|---|
| | NCT05227664 | AK112, Nab Paclitaxel, and Paclitaxel | Metastatic TNBC and Locally Advanced TNBC | II | 120 | Recruiting |
| | NCT05382442 | AK117, Oxaliplatin, Capecitabine, Irinotecan, Leucovorin, and 5-fluorouracil | Metastatic Colorectal Cancer | II | 254 | Active |
| | NCT06196203 | Azacitidine | High-risk Myelodysplastic Syndromes | II | 90 | Recruiting |
| | NCT04728334 | NA | Malignant Neoplasms | I | 49 | Completed |
| | NCT06642792 | AK129 | Classic Hodgkin's Lymphoma | I/II | 280 | Recruiting |
| Magrolimab (Hu5F9-G4) | NCT02678338 | NA | Hematologic Neoplasms | I | 20 | Completed |
| | NCT02216409 | NA | Solid Tumor | I | 88 | Completed |
| | NCT03248479 | Azacitidine | Hematologic Neoplasms | Ib | 258 | Terminated |
| | NCT05807126 | NA | Breast Cancers and Prostate Cancers | I | 0 | Withdrawn |
| | NCT04599634 | Obinutuzumab, Venetoclax, Acetaminophen, Diphenhydramine, Prednisone/prednisolone, and Methylprednisolone | Follicular Lymphoma, Marginal Zone Lymphoma, Mantle Cell Lymphoma Chronic Lymphocytic Lymphoma, and B-cell Lymphoma | I | 11 | Completed |
| | NCT03527147 | AZD9150, Acalabrutinib, AZD6738, Rituximab, and AZD5153 | Non-hodgkin's Lymphoma and Diffuse Large B-cell Lymphoma | I | 30 | Completed |
| | NCT05367401 | Sabatolimab, Magrolimab, and Azacitidine | Myelodysplastic Syndromes and Acute Myeloid Leukemia | Ib/II | 0 | Withdrawn |
| | NCT04788043 | Pembrolizumab | Classic Hodgkin's Lymphomas | II | 8 | Active |
| | NCT05169944 | NA | Brain Cancers and Pediatric Brain Tumor | I | 13 | Completed |
| Gentulizumab | NCT05221385 | NA | Solid Tumor and Non-Hodgkin's Lymphoma | I | 58 | Terminated |
| | NCT05263271 | NA | Acute Myelogenous Leukemia and Myelodysplastic Syndromes | I | 58 | Unknown |
| STI-6643 | NCT04900519 | NA | Solid Tumors | I | 100 | Completed |
| TQB2928 | NCT05192512 | NA | Advanced Cancers | I | 180 | Unknown |
| | NCT06438783 | Anlotinib | Osteosarcoma and other Solid Tumors | I | 43 | Recruiting |
| | NCT06297642 | Penpulimab | Advanced Malignant Neoplasm | I | 3 | Terminated |
| | NCT06008405 | NA | Acute Myeloid Leukemia Myelodysplastic Syndromes | I | 48 | Unknown |
| | NCT06585059 | Almonertinib Mesilate tablets | Advanced Non-small Cell Lung Cancer | Ib | 20 | Not Yet Recruiting |

| | | | | | | | |
|---|---|---|---|---|---|---|---|
| | | NCT04854681 | NA | Advanced Solid Tumors and Hematological Malignancies | I | | Unknown |
| *Monoclonal anti-CD47* | CC-90002 | NCT02367196 | Rituximab | Hematologic Neoplasms | I | 60 | Completed |
| | | NCT02641002 | NA | Myeloid Leukemia and Acute Myelodysplastic Syndromes | I | 28 | Terminated |
| | HMPL-A83 | NCT05429008 | NA | Advanced Tumors | I | 40 | Completed |
| | IMC-002 | NCT05276310 | NA | Advanced Cancer | I | 62 | Recruiting |
| | SRF231 | NCT04306224 | NA | Solid Tumor and Lymphomas | I | 12 | Completed |
| | | NCT03512340 | NA | Advanced Solid Cancers and Hematologic Cancers | I/Ib | 148 | Completed |
| | Not Specified | NCT04588324 | SHR2150 | Solid Tumor | I/II | 50 | Unknown |
| | Not Specified | NCT05266274 | Azacitidine | Recurrent Acute Myelogenous Leukemia After Transplantation | Not Specified | 69 | Completed |
| *Fusion protein (SIRPα IgG1 Fc)* | Evorpacept (ALX148) | NCT05429008 | Pembrolizumab and Doxorubicin | Ovarian Cancer | II | 31 | Recruiting |
| | | NCT03013218 | Pembrolizumab, Trastuzumab, Rituximab, and Ramucirumab + Paclitaxel, 5-Fluoruoracil + Cisplatin | Metastatic Cancer, Solid Tumor, and Non-Hodgkin's Lymphoma | I | 174 | Active |
| | | NCT07007559 | Cetuximab, FOLFIRI (Irinotecan, 5-Fluorouracil, and Leucovorin), Trastuzumab, Paclitaxel, Capecitabine, Eribulin, Gemcitabine, and Vinorelbine | Breast Cancer and Metastatic Colorectal Cancer | Ib/II | 280 | Recruiting |
| | | NCT04755244 | Venetoclax and azacitidine | Acute Myeloid Leukemia | I | 14 | Terminated |
| | | NCT05167409 | Cetuximab and Pembrolizumab | Microsatellite Stable Metastatic Colorectal Cancer | II | 48 | Active |
| | | NCT04675294 | Pembrolizumab | Head and Neck Cancer and Head and Neck Squamous Cell Carcinoma | II | 189 | Active |
| | | NCT05002127 | Trastuzumab, Ramucirumab and Paclitaxel | Gastric Cancer, Gastroesophageal Junction Adenocarcinoma, and Gastric Adenocarcinoma | II/III | 450 | Active |
| | | NCT04675333 | Pembrolizumab, Cisplatin/Carboplatin, and 5-Fluoruoracil | Head and Neck Cancer and Head and Neck Squamous Cell Carcinoma | II | 172 | Active |
| | | NCT04417517 | Azacitidine | Higher Risk Myelodysplastic Syndromes | I/II | 65 | Active |
| | | NCT05025800 | Lenalidomide and Rituximab | Non-Hodgkin Lymphomas (B-Cell and Composite LymphomaIndolent B-Cell), B-Cell Lymphomas (B-Cell and Mediastinal), Lymphomas (Follicular, Mantel Cell, Marginal Zone, Mediastinal, and Ann Arbor), | I/II | 47 | Active |

| | | | | | | | |
|---|---|---|---|---|---|---|---|
| | | | | Transformed to Diffuse Large B-Cell Lymphomas, and Richter Syndrome | | | |
| | | NCT05524545 | Enfortumab Vedotin | Bladder Cancer and Urothelial Carcinoma | I | 36 | Complete |
| | | NCT05467670 | Pembrolizumab and Doxorubicin | Ovarian Cancer | II | 31 | Recruiting |
| | | NCT05868226 | Fam-Trastuzumab Deruxtecan-Nxki, Zanidatamab, and Tucatinib | HER2-positive Breast Cancers, HER2-low Breast Cancers, HER2-negative Breast Cancer, TNBC, Estrogen Receptor Positive Tumors, Progesterone Receptor-positive Breast Cancers, and Solid Tumors | I/IIb | 124 | Recruiting |
| | | NCT04643002 | Isatuximab, Dexamethasone, Pomalidomide, Belantamab mafodotin, Pegenzileukin, SAR439459, and Belumosudil | Plasma Cell Myeloma Refractory | I/II | 258 | Recruiting |
| | Ontorpacept (TTI-621) | NCT02663518 | Rituximab and Nivolumab | Hematologic Malignancies and Solid Tumor | I | 249 | Terminated |
| | | NCT04996004 | Doxorubicin | Leiomyosarcoma | II | 76 | Terminated |
| | | NCT02890368 | PD-1/PD-L1 Inhibitor, pegylated interferon-α2a, Talimogene laherparepvec, and radiation | Solid Tumors, Mycosis Fungoides, Melanoma, Merkel Cell Carcinoma, Breast Cancer, HPV Related Neoplasm, and Soft Tissue Sarcoma | I | 56 | Terminated |
| *Fusion protein (SIRPα IgG4 Fc)* | Maplirpacept (TTI-622) | NCT05626322 | Tafasitamab, Lenalidomide | Diffuse Large B-Cell Lymphoma | Ib/II | 6 | Terminated |
| | | NCT05139225 | Daratumumab hyaluronidase-fihj | Multiple Myeloma | I | 7 | Active |
| | | NCT05261490 | Pegylated Liposomal Doxorubicin | Ovarian Cancer, Fallopian Tube Cancer, Epithelial Ovarian Cancer, and Primary Peritoneal Carcinoma | I/II | 10 | Terminated |
| | | NCT05896163 | Glofitamab and Obinutuzumab | Diffuse Large B-Cell Lymphoma | Ib/II | 70 | Recruiting |
| | | NCT03530683 | Azacitidine, Venetoclax, Carfilzomib, Dexamethasone, Anti-CD20, and Isatuximab | Lymphoma, Multiple Myeloma, Acute Myeloid Leukemia, and Diffuse Large B-Cell Lymphoma | I | 189 | Terminated |
| | | NCT05507541 | Ontorpacept and Pembrolizumab | Diffuse Large B-Cell Lymphomas, Large B-Cell Lymphomas, Follicular Lymphomas, and Gray Zone Lymphomas | II | 10 | Active |
| | | NCT05896774 | NA | Non-Hodgkin Lymphoma and Multiple Myeloma | I | 10 | Completed |
| | | NCT05567887 | NA | Lymphoma and Multiple Myeloma | I | 7 | Completed |
| | | NCT05675449 | Elranatamab and Carfilzomib | Multiple Myeloma | Ib | 90 | Recruiting |
| | HCB101 | NCT05892718 | NA | Advanced Solid Tumor and Refractory Non-Hodgkin's Lymphoma | I | 60 | Recruiting |

| | | | | | | | |
|---|---|---|---|---|---|---|---|
| | | NCT06771622 | Trastuzumab, Pertuzumab, Oxaliplatin, Capecitabine, Ramucirumab, Paclitaxel, Bevacizumab, Cetuximab, Irinotecan, Leucovorin, 5-FU, Toripalimab, Albumin-bound paclitaxel, Pembrolizumab, and Gemcitabine | Solid Cancers | I/II | 150 | Recruiting |
| | | NCT07204574 | Bevacizumab, Cetuximab, FOLFIRI, and FOLFOX | Colorectal Cancers | I/II | 40 | Recruiting |
| | | NCT07136545 | Pembrolizumab | Head and Neck Squamous Cell Carcinoma | I/II | 50 | Recruiting |
| *Small molecule* | AUR103 | NCT05607199 | Azacitidine | Solid Tumor, Acute Myeloid Leukemia, Myelodysplastic Syndromes, and Non-Hodgkin's Lymphoma | I | 80 | Recruiting |
| | | NCT07040059 | Trastuzumab + CAPOX | HER2 Positive Gastric Cancer and Gastroesophageal Junction Adenocarcinoma | I/IIb | 18 | Recruiting |

**Supplementary Table 2** SIRPα targeted agents in clinical trials for cancers

| ***Drug Format*** | ***Drug Name/Code*** | ***Clinical trial ID*** | ***Co-therapies*** | ***Cancer Types*** | ***Phase*** | ***N*** | ***Trial Status*** |
|---|---|---|---|---|---|---|---|
| *Monoclonal anti-SIRPα* | BYON4228 | NCT06932952 | Pembrolizumab | Solid Tumor | I | 85 | Not Yet Recruiting |
| | | NCT05737628 | Rituximab | Lymphoma | I | 85 | Recruiting |
| | DS-1103a | NCT05765851 | Trastuzumab derextecan and Enhertu®/ T-DXd | Advanced Solid Tumor and Breast Cancer | I | 85 | Active |
